# Reduced Permeability to Rifampicin by Capsular Thickening as a Mechanism of Antibiotic Persistence in *Mycobacterium tuberculosis*

**DOI:** 10.1101/624569

**Authors:** Jees Sebastian, Sharmada Swaminath, Parthasarathi Ajitkumar

**Author notes:** Address correspondence to P. Ajitkumar,.

## Abstract

Persisters constitute a subpopulation of bacteria that can tolerate lethal concentrations of antibiotics. Multiple mechanisms have been suggested for bacterial persistence against antibiotics. With mycobacteria being no exception to this behaviour, we had reported the *de novo* emergence of genetically antibiotic-resistant *Mycobacterium tuberculosis* from persister cells upon prolonged exposure to microbicidal concentrations of the anti-tuberculosis drugs, rifampicin and moxifloxacin. Here, we present evidence for reduced permeability to rifampicin as a mechanism for persistence of *Mycobacterium tuberculosis in vitro*. We observed that rifampicin persistent *M. tuberculosis* cells developed a thick outer layer (TOL) capsule. The TOL restricted the entry of fluorochrome-conjugated rifampicin, 5-carboxyfluorescein-rifampicin (5-FAM-rifampicin), which retained only 2.5% of its original bactericidal activity, but high levels of permeability, on actively growing mid-log phase cells. Gentle mechanical removal of TOL significantly enhanced 5-FAM-rifampicin entry into the persister cells. The level of 5-FAM-rifampicin in the persister cells was not affected by the pre-incubation of the cells with verapamil, a drug efflux pump inhibitor, ruling out the involvement of efflux pumps in the reduced intracellular concentration of 5-FAM-rifampicin. GC-MS analysis of TOL showed the presence of ∼7-fold, ∼5-fold and ∼2- fold higher levels of α-D-glucopyranoside, 1,2,5-linked-mannitol, and 3,4-linked mannose, respectively, among ∼2-fold higher levels of derivatives of several other types of sugars such as arabinose and galactose. Taken together, the present study reveals that rifampicin-persistent *M. tuberculosis* cells develop TOL that enables the bacilli to restrict entry of rifampicin and thereby remain tolerant to the antibiotic *in vitro*.

## INTRODUCTION

*Mycobacterium tuberculosis*, which is the causative agent of tuberculosis, is one of the most successful human pathogens due to its ability to survive under diverse extreme stress conditions. Like many other pathogenic bacteria, *M. tuberculosis* also exhibits several survival strategies to overcome the lethality imposed by antibiotic exposure. One of the unique features of *M. tuberculosis* that has enabled the bacilli to remain refractory to many commonly used antibiotics is the distinct cell wall structure (1). The complex cell wall of *M. tuberculosis* is well known for imposing limited permeability to several host-derived antimicrobial biomolecules that makes the bacilli tolerant to many toxic components from its normal habitats (2-5). Studies have been conducted on the physiological significance of the changes that the cell wall undergo inside macrophages (6), under extreme nutritional stress conditions (7) and hypoxia (8). However, there is no information on whether the cell envelope has any contribution to the persistence of the bacilli against antibiotics.

Persistence is one of the survival strategies of mycobacteria to remain tolerant to lethal concentrations of antibiotics. Although several mechanisms that contribute to mycobacterial persistence have been reported (9, 10), very little information is available on the ultrastructural features of mycobacterial antibiotic persisters and their contribution, if any, to antibiotic tolerance. Earlier studies in *Escherichia coli* and *Staphylococcus aureus* have shown that exposure to a sub-lethal concentration of antibiotics can cause the emergence of resisters through the generation of reactive oxygen species and consequential mutagenesis (11). Subsequently, in our study, *M. tuberculosis* persisters were found in the presence of lethal concentrations of rifampicin and moxifloxacin, leading to the *de novo* emergence of antibiotic resisters (12). These two studies raised the possibility that the bacilli might have some hitherto unknown strategy to ensure sub-lethal concentration of antibiotics inside persister cells even though they remain exposed to lethal concentrations of the antibiotics. In order to verify this possibility, we examined whether the permeability of rifampicin into *M. tuberculosis* persister cells was affected by any ultrastructural changes. The present study reports the ultrastructural changes of rifampicin persistent *M. tuberculosis* cells and their contribution to the restricted permeability of rifampicin, in comparison with the entry of rifampicin into actively growing mid-log phase cells.

## MATERIALS AND METHODS

### Rifampicin treatment of *Mycobacterium tuberculosis*

*Mycobacterium tuberculosis* H_37_R_a_ (National JALMA Institute of Leprosy and Other Mycobacterial Diseases, Agra, India) was used in all the experiments. Bacteria were grown in Middlebrook 7H9 broth (BD) supplemented with 0.2% glycerol, 0.05% Tween-80 and 10% ADS (albumin, dextrose, sodium chloride) until mid-log phase (MLP, 0.6 at OD_600 nm_) at 37°C under shaking at 170 rpm. Cultures were then exposed to 1 µg/ml (10x MBC; 12) of freshly prepared rifampicin (Sigma) under the same condition. Aliquots of the culture were withdrawn once in every 24 hrs and used for serial dilution and plating on sterility checked Middlebrook 7H10 agar supplemented with 10% ADS in the absence of rifampicin. Plates were sealed with parafilm and incubated at 37°C in CO_2_ incubator (5% CO_2_) for determining CFU. For experiments using 5-FAM-rifampicin (Merck-Millipore), MLP or persistence phase cells were exposed to 1.5 µg/ml (concentration equimolar to 10x MBC rifampicin) for the required period and taken for flow cytometry analysis (see below).

### Transmission electron microscopy

M. *tuberculosis* cells from different stages of rifampicin treatment were harvested and processed for transmission electron microscopy as previously described (13). In brief, the cells were prefixed in 1% (w/v) osmium tetroxide (solution in double-distilled water) in 0.15 M cacodylate buffer (pH 7.2) for 1 hr at room temperature and washed once in the same buffer. Further, cells were fixed using 2% (v/v) glutaraldehyde and 2% (w/v) tannic acid in 0.15 M cacodylate buffer for 2 hrs at room temperature. Cells were washed once in the same buffer and re-fixed in 1% osmium tetroxide overnight at 4°C. Samples were dehydrated by a series of ethanol washes (with 25%, 50%, 75% and 95% ethanol) (5 min incubation between every wash) and infiltrated with 50% LR white resin in 50% ethanol for 24 hrs at 4°C. Dehydrated cells were collected by centrifugation and the pellet was used for making blocks in gelatine capsules with 100% LR white resin. Blocks were solidified by incubating at 65°C for 2-3 days in a dry bath. Ultrathin sections of 80 to 100 nm were prepared from the blocks using ultramicrotome and collected on copper grids. Sections were stained with uranyl acetate (0.5%) and lead citrate (0.04%) and observed under JEOL-100 CX II electron microscope at 100 kV.

For ruthenium red staining, we followed the previously reported method (14). In brief, the cells were harvested and washed once in Middlebrook 7H9 medium and prefixed in the presence of paraformaldehyde and glutaraldehyde (2.5% w/v each), CaCl_2_ and MgCl_2_ (5 mM each) and 0.05% (w/v) ruthenium red in 0.1 M cacodylate buffer (pH 7.2) for 2 hrs at 4°C. Cells were washed once in the same buffer and incubated with 2.5% w/v glutaraldehyde and 0.05% (w/v) of ruthenium red in cacodylate buffer. Cells were washed once and post-fixed in osmium tetroxide (OsO_4_) and uranyl acetate (1% w/v of each) containing ruthenium red (0.05% w/v). Fixed cells were dehydrated with a series of ethanol washes (with 25%, 50%, 75% and 95% ethanol) (5 min incubation between every wash, ethanol contained 0.05% w/v of ruthenium red) and infiltrated with 50% LR white resin in 50% ethanol for 24 hrs at 4°C. Cells were collected by centrifugation and the pellet was used for making blocks in gelatine capsules with 100% LR White resin. Blocks were solidified by incubating at 65°C for 2-3 days in dry bath. Ultrathin sections of 80-100 nm were prepared from the blocks using ultra-microtome and placed on copper grids. Sections were stained with uranyl acetate (0.5%) and lead citrate (0.04%) and observed under JEOL-100 CX II electron microscope at 100 kV.

### Bacterial hydrophobicity assay

A modified protocol of a previously reported method (15, 16) was used. Cells from MLP and rifampicin persistence phase were harvested by centrifugation and resuspended in filter sterilised PUM buffer (100 mM K_2_HPO_4_.3H_2_O, 50 mM KH_2_PO_4_, 33.3 mM urea, 1 mM MgSO4.7H_2_O, in double-distilled autoclaved water, pH 7.1) to get an approximate density of 0.7 at OD_600 nm_. Phase extraction was performed in siliconised borosilicate tubes using one volume of PUM buffer containing cells against three volumes of n-hexadecane and vortexed for 8-10 seconds and left at room temperature for 15 min. The aqueous phase was collected into siliconised microcentrifuge tubes using siliconised tips, mildly sonicated and used for serial dilution and plating on sterility checked Middlebrook 7H10 agar plates containing 10% ADS supplement. The difference in the CFU of the aqueous phase before and after phase separation was used for calculating the percentage of cells with hydrophilic outer layer.

### Estimation of zeta potential of mycobacterial persister cells

For the determination of surface charge of *M. tuberculosis* cells, aliquots of the culture were withdrawn on different days during rifampicin treatment and washed once with Middlebrook 7H9 broth. Cells were resuspended in fresh medium and used for zeta potential measurement using zeta sizer nano series (Nano-ZS90, Malvern Instruments). For the estimation of the isoelectric point of the cells, the cells were resuspended in solutions of varying pH from 2 to 10 in PPMS buffer (40 mM of K_2_HPO_4_, 20 mM of KH_2_PO_4_, and 1.5 mM of MgSO_4_.7H_2_O per litre of Milli Q water, pH adjusted using HCl or NaOH). Zeta potential values of the cells at different pH were calculated and the graph was plotted, as described (17, 18).

### GC-MS analysis of OL

Cells from MLP and persistence phase were collected and used for OL extraction as previously reported (19) with minor modifications. Briefly, cells were washed once in 1x PBS and resuspended in 20 ml distilled water followed by incubation with 10 gm of sterile 4 mm glass beads at 50 rpm for 15 min. The cell suspension was collected and centrifuged at 12000 xg for 10 min. The supernatant was filtered through a 0.2 micron filter and the filtrate was lyophilized. The sample was derivatised and subjected to GC-MS performed at Mass spectrometry glycomics facility at C-CAMP Bangalore. Data analysis was also performed at the same facility.

### Fluorescence microscopy

Cells from MLP and persistence phases were harvested and resuspended in 100 µl of Middlebrook 7H9 broth. For the staining of cells, 5-FAM-rifampicin (1.5 µg/ml) or calcofluor white (CFW) (1:1000 dilutions of 0.1% solution, Sigma) was added into the cell suspension and incubated for 1 hr at 37°C incubator shaker. Propidium iodide (1:1000) was added (to discriminate dead cells) and layered over poly-L-lysine treated multiwell slides 20 min in the dark. Wells were washed once with phosphate buffered saline (PBS) and mounted with glycerol and cover slips and observed under 100X objective in Zeiss AxioVision fluorescence microscope. Bead beating was performed by incubating 20 ml culture with 10 grams of glass beads (4 mm) at 37°C for 15 min at 50 rpm, prior to the addition of 5-FAM-rifampicin and processed similarly as above.

### Flow cytometry analysis

Cell suspension (500 µl) was exposed to 1.5 µg/ml of 5-FAM-rifampicin and incubated at 37°C incubator for 1 hr in the dark. Aliquots were collected at specified intervals and cells were harvested by centrifugation at 12000 x g for 5 min at 4°C and washed once with ice-cold Middlebrook 7H9 broth and used for flow cytometry analysis. For bead beating, 20 ml culture was incubated with 10 grams of glass beads (4 mm) at 37°C for 15 min at 50 rpm prior to the addition of 5-FAM-rifampicin and processed similarly for flow cytometry with 488 nm solid state laser and 527/32 nm emission filter (for GFP) at low or medium flow rate. For relative permeability estimation of rifampicin persisters for 5-FAM-rifampicin, cells from MLP, persistence phase and verapamil-pretreated (50 µg/ml) persisters were incubated with 1.5 µg/ml of 5-FAM-rifampicin and incubated for 1 hr at 37°C in the dark and processed in the same way as described earlier.

For CFW staining and OL analysis, 500 µl aliquots of the cells before and after bead beating were incubated with 1:1000 dilutions of CFW (0.1% solution, Sigma) for 1 hr at 37°C. Cells were washed once with Middlebrook 7H9 broth and used for flow cytometry analysis in BD FACSVerse flow cytometer.

About 10000 gated cells were analysed and used for the estimation of the median fluorescence value of the 5-FAM-rifampicin stained cells while keeping the autofluorescence at 2-log_10_ value. The photomultiplier tube (PMT) voltage settings used for measuring the 5-FAM-rifampicin fluorescence of the cells were 208 (FSC), 333 (SSC). The instrument was calibrated using FACSuite cytometer set up and tracking beads (CS&T, Becton Dickinson). Data were processed and analysed using FACSuite software and the statistical significance between the time points was calculated using paired t-test of GraphPad Prism version 5.0.

### 5-FAM-rifampicin bioassay

Rifampicin sensitive *Staphylococcus aureus* (ATCC 25923) was used for the bioassay of 5-FAM-rifampicin (20). In brief, rifampicin (sigma) and 5-FAM-rifampicin were dissolved in DMSO to make a stock solution of 2 mg/ml. Dilutions were made from the stock solution and used for the agar diffusion assay. LB agar plates were made by mixing 50 µl of the *Staphylococcus aureus* glycerol stock with 100 ml LB agar (warm to the touch). Wells were made in the agar using a stainless steel puncture with 0.5 cm diameter. Known concentrations of rifampicin or 5-FAM-rifampicin solutions (50 µl) were added into the well and incubated overnight at 37°C. The diameter of the zone of inhibition was measured using Vernier caliper and used for plotting the standard graph for rifampicin, from which and the zone of inhibition of 5-FAM-rifampicin, the bioactivity of 5-FAM-rifampicin was calculated.

### 5-FAM-rifampicin permeability assay

To estimate the extent of 5-FAM-rifampicin entry into MLP cells and rifampicin persisters, cultures were treated with a final concentration of 1.5 µg/ml of the drug conjugate in 20 ml culture and incubated in a shaker at 37°C. Cultures for both (MLP and persisters) were mildly beaten with 4 mm glass beads to remove the outer capsular layer, as a control for the experiment. Aliquots were taken at every 15 min intervals and quickly washed once with ice-cold Middlebrook 7H9 broth and used for flow cytometry analysis using BD FACSVerse system. Samples before 5-FAM-rifampicin addition (0 min) were used as the control for autofluorescence and median of fluorescence was kept at 10^2^.

To construct a standard graph for 5-FAM-rifampicin entry, MLP cells were incubated with two-fold increasing concentrations of 5-FAM-rifampicin in an incubator shaker for 1 hr at 37°C. Cells were harvested and quickly washed once with ice-cold Middlebrook 7H9 broth and used for flow cytometry analysis in BD FACSVerse system. Autofluorescence median of the untreated cells was kept at 10^2^ and the normalised median values were used for plotting standard graph. Rifampicin persisters, which were isolated from the 12^th^ day of the treatment, were incubated with 5-FAM-rifampicin equivalent of 10x MBC rifampicin. These cells were also similarly processed and readings were taken from 10000 cells. Normalised fluorescence values from six independent persister cell samples were used for calculating the relative concentration of 5-FAM-rifampicin in persister cells from the standard graph.

### RESULTS AND DISCUSSION

Earlier studies from our laboratory had shown that actively growing MLP *M. tuberculosis* cells exposed to 10x MBC concentration of rifampicin consistently showed a triphasic response involving killing phase (0 to 10^th^ day of exposure), persistence phase (10^th^ to 15^th^ day of exposure), followed by regrowth phase (beyond 15^th^ day of exposure) (12). As part of our earlier characterisation of the triphasic response of *M. tuberculosis* cells to 10x MBC rifampicin, we had found that the rifampicin concentration stays at ∼5x MBC even on the 15^th^ day of the prolonged exposure that spanned for 20 days (12). Therefore, for the present ultrastructural studies, the persistence phase cells from the 12^th^ day of exposure to 10x MBC rifampicin were used.

### Rifampicin persister cells possess thick outer layer (TOL)

Transmission electron micrographs of rifampicin persister *M. tuberculosis* cells, stained with tannic acid-lead citrate combination (13), showed a significantly thick, but strikingly uneven, loosely bound and deeply stained outer layer (OL) (**Fig. 1A**). On the contrary, the MLP cells (control) showed an evenly thin outer layer, which we called normal outer layer (NOL) (**Fig. 1B**). The TOL thickness of persister cells ranged between 25-130 nm (n = 27), while the MLP cells showed a more-or-less evenly thin layer of NOL with an average thickness of ∼20 nm (**Fig. 1C**). However, the thickness of electron transparent layer (ETL) of the persister and the control cells were found comparable and were morphologically like the already published data (**Fig. 1D**) (14, 21, 22). Since rifampicin exposure for the experimental sample was started from the MLP cells, cultures from MLP stage for an equivalent number of days, but unexposed to rifampicin did not show any TOL (**Fig. S1**). Thus, the ultrastructural difference between the rifampicin persister cells and the control cells was confined to the outer layer (OL) thickness.

**FIG 1.**
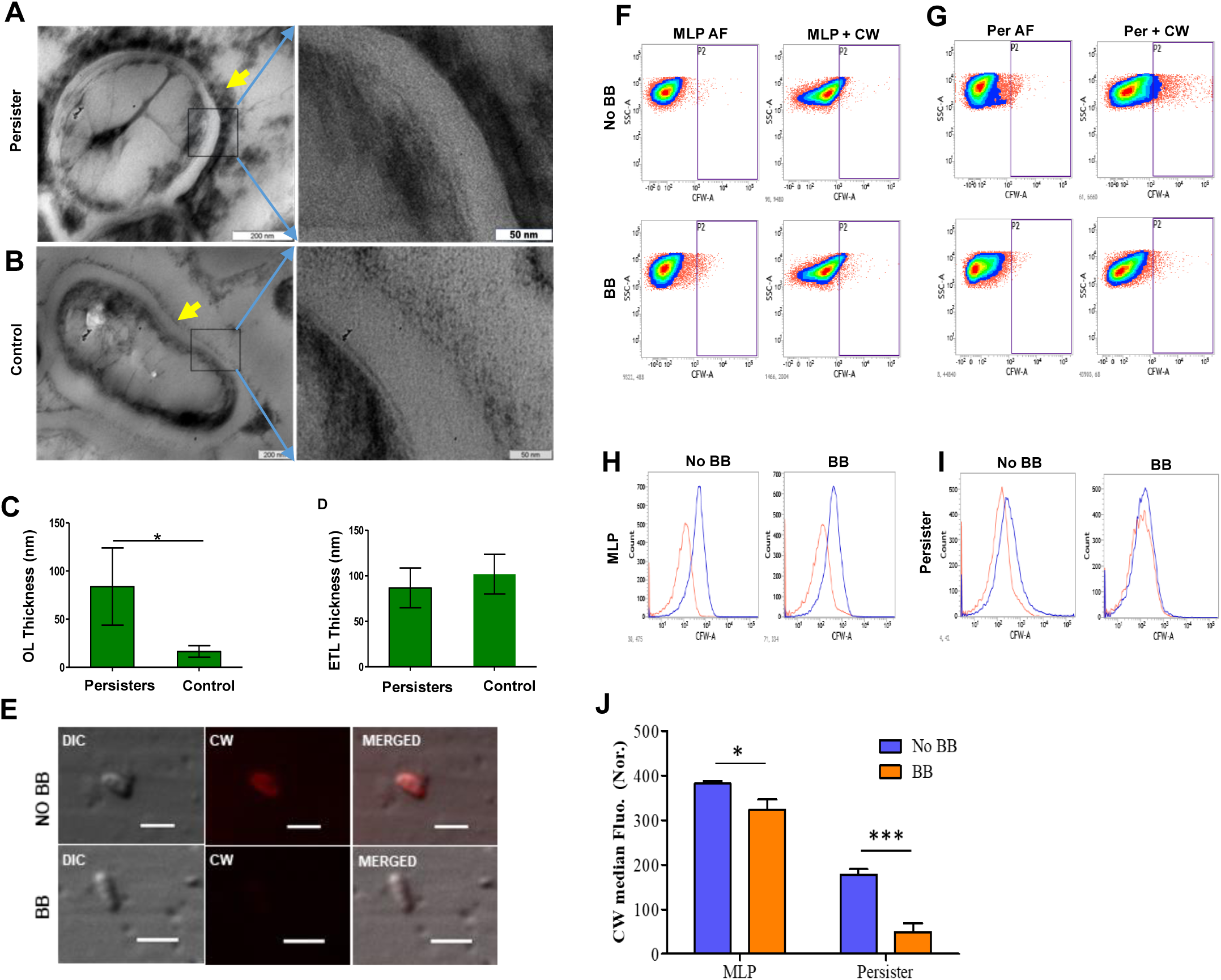
Presence of TOL in rifampicin persister cells. Transmission electron micrograph of *M. tuberculosis* cells showing differential thickening of OL in persister cell (**A**) and in control MLP cells (**B**). Magnified images are shown right side, the yellow arrows indicate OL. Thickness (in nm) of OL (**C**) and ETL (**D**) between persister and MLP cells. (**E**) CFW staining of OL in persister cells before and after bead beating (BB). Scale, 2 µm. Density plots of the flow cytometry analysed CFW stained MLP cells (**F**) and persister cells (**G**) before and after BB. AF, autofluorescence. Histogram overlay of the flow cytometry analysed CFW stained MLP cells (**H**) and persister cells (**I**) before and after BB. (**J**) Bar graph showing normalised median fluorescence intensity of CFW in MLP and persister cells before and after BB. One asterisk (*) indicates P value less than or equal to 0.05 (P ≤ 0.05). Three asterisks (***) indicate P value lesser than 0.001 (P < 0.001). The statistical significance was calculated using two-tailed paired t-test.

Transmission electron microscopy of persister and MLP cells stained with ruthenium red was performed to specifically stain polysaccharides for detection (14). It showed the presence of polysaccharides as an uneven layer on the surface of rifampicin persister cells unlike a very thin layer on the MLP control cells (n = 22) (**Fig. S2**). The thin OL on the MLP also contained polysaccharides, as reported (19). The presence of polysaccharides was further confirmed by staining with a polysaccharide specific fluorophore, calcofluor white (CFW; 23, 24) (**Fig. 1E, top panel**). The persister cells, which were gently bead beaten to remove OL, as described (19), and stained with CFW, showed loss of CFW fluorescence (**Fig. 1E, bottom panel**). Flow cytometry analysis of CFW stained native and bead beaten persister cells also showed loss of CFW fluorescence. However, the MLP cells (**Fig. 1F, H, J**) did not show an appreciable difference in CFW fluorescence between the native and the bead beaten samples, compared to persister cells (**Fig. 1G, I, J**). Thus, by using multiple staining methods with two different polysaccharide specific reagents confirmed the differential polysaccharide content on the MLP and persister cells.

### TOL imparts hydrophilicity to rifampicin persister cells’ surface

It was shown that the constituents of the OL of actively growing mycobacteria are mostly polysaccharides and proteins with low lipid content (19). Therefore, due to the thickening of the OL in persister cells, the possibility of a change in the hydrophilicity of the surface of persister cells, as compared to that of the actively growing MLP cells, was verified. For this purpose, cells from the rifampicin persistence phase and MLP were subjected to phase separation between highly hydrophobic hexadecane and aqueous buffer to measure cell-surface hydrophobicity, as described (15). The cells retained in the aqueous phase after the phase extraction reflects the proportion of cells that have a hydrophilic surface. We observed that an average 7% of the persister population was hydrophilic and retained in the aqueous phase, while only 0.03% of MLP cells being hydrophilic (**Fig. 2A**). This showed that the persister population contained a relatively higher proportion of cells having hydrophilic surface.

**FIG 2.**
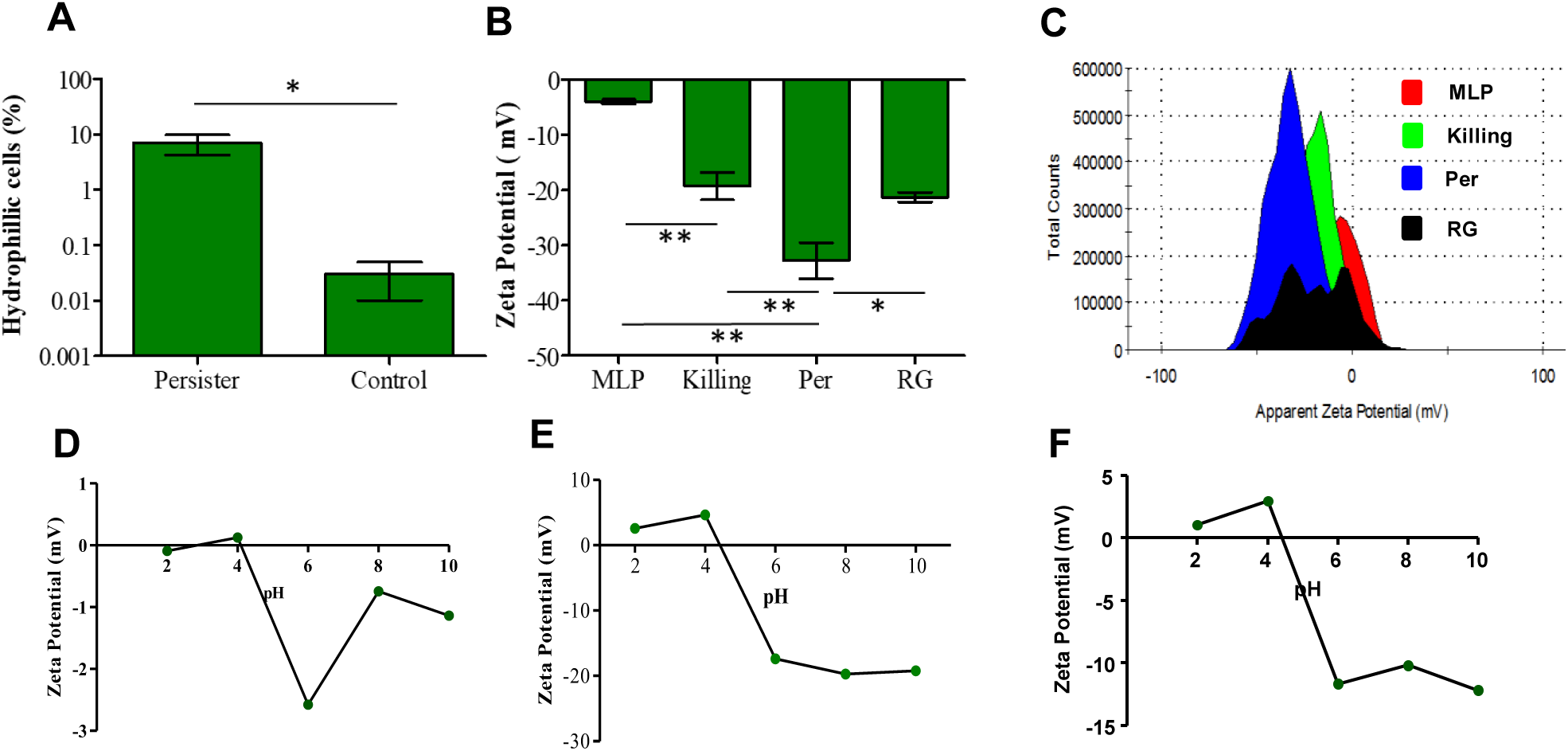
Physicochemical properties of rifampicin-exposed *M. tuberculosis* persister cells. (**A**) Hexadecane assay showing the proportions of hydrophilic rifampicin persister cells and MLP cells. (**B**) ZP of cells from different phases of rifampicin exposed of *M. tuberculosis* cells and its corresponding (**C**) histogram overlay. ZP of TOL as a function of pH showing the isoelectric point of the TOL between pH 4.0 and pH 5.0 for: (**D**) MLP cells, (**E**) persister cells, and (**F**) cells from killing phase. One asterisk (*) indicates P value less than or equal to 0.05 (P ≤ 0.05). Two asterisks (**) indicate P value less than or equal to 0.01 (P ≤ 0.01). The statistical significance was calculated using two-tailed paired t-test.

### Rifampicin persister *M. tuberculosis* cells possess negative surface charge

The presence of anionic polysaccharides in the TOL could impart a negative surface charge to the persister cells and confer hydrophilicity. The cell surface negative charge can be measured by the zeta potential (ZP) of intact cells (17, 18). The ZP of *M. tuberculosis* cells was measured for MLP cells and the cells from the killing, persistence and regrowth phases of rifampicin exposure, which were described before (12). *M. tuberculosis* MLP cells showed a ZP value of -3.91 mV, with a gradual increase in the negative surface charge over the course of rifampicin exposure (**Fig. 2B**). The gradual increase in the negative ZP value upon continued exposure to rifampicin indicated accumulation of negatively charged molecules on the cell surface. Cells from the killing phase showed a ZP of -19.26 mV indicating a remarkable increase in their negative surface charge compared to that of the rifampicin-unexposed MLP cells. The persistence phase cells showed a peak ZP value of -32.76 mV, showing that the persister cells have significantly high negative surface charge density compared to that of the cells from any other phase of rifampicin exposure. The high negative ZP of persister cells indicated the accumulation of anionic polysaccharides on the cell surface. The regrowth phase cells showed a reduction in the negative ZP value indicating the loss of TOL once the cells gained rifampicin resistance and came out of the persistence phase. In addition, we observed multiple peaks in the ZP histogram of RG phase population, suggesting the possibility of heterogeneity in terms of surface charge among the rifampicin-tolerant/resistant regrowing cells (**Fig. 2C**).

In order to determine ionic properties of the mycobacterial TOL, the isoelectric point (pI) of *M. tuberculosis* cells was measured at different pH values (2, 4, 6, 8 and 10). The pH at which cells show zero ZP value was considered as the pI of the cells. To estimate the isoelectric point (pI) of *M. tuberculosis* cells, at different pH values, we measured the ZP of cells from various time points during rifampicin exposure. The positive ZP of MLP cells at pH 2.0 and pH 4.0 were dropped to -2.58 at pH 6.0 indicating that the pI of MLP cells were between pH 4.0 and pH 6.0 (**Fig. 2D**). The persistence phase cells showed a similar pI profile with a lower ZP value at high pH (**Fig. 2E**). Although the cells in the killing phase also showed a similar pI, the extent of negative ZP potential at high pH was considerably lesser than that of the persistence phase cells (**Fig. 2F**). Thus, *M. tuberculosis* cells from MLP, killing and persistence phases showed similar pI values with a varying surface charge at higher pH, probably due to the high levels of anionic polysaccharide content on their surface. The comparable pI values of the MLP cells and persisters indicated that the ionic properties of the TOL might not have changed over the period of rifampicin exposure. This alluded to the possibility that the nature of the molecules might be similar but their relative composition might be different (see GCMS data described below), thereby having similar pI.

### Molecular analysis of TOL composition

A comparative molecular analysis using GC-MS was performed to find out the polysaccharide composition of TOL on persistence phase cells and its difference from that of the NOL on MLP cells. For this purpose, the NOL from MLP cells and TOL from persister cells were gently extracted, as described (19) and used for mass spectrometry. The percentage of relative abundance of the monosaccharides detected are listed in **Table 1**. We observed ∼5-fold increase in the levels of 1, 2, 5-mannitol and ∼6-7-fold increase in the α-D-glucopyranoside levels. This composition is consistent with an earlier study that the NOL of actively growing *M. tuberculosis* (19). The thickening of the OL has caused several fold increase in the levels of the constituents that were present in the NOL of MLP cells (see **Table 1**). For instance, the 6-7-fold higher levels of α-D-glucopyranoside, as a breakdown product in the GC-MS analysis, alluded to the possibility of the presence of trehalose, which is α-D-glucopyranosyl-α-D-glucopyranoside (glucose disaccharide), known to desiccate bacterial cells against severe stress conditions (25-27). Similarly, glucan, which is a component of the *M. tuberculosis* cell surface, is a polymer of glucose that is expressed *in vitro* and *in vivo* (28). The presence of high levels of α-D-glucopyranoside might be an indication of the presence of glucan as well. The increased levels of arabinose and mannose probably signify the presence of arabinomannan, which is one of the significant components of OL (1, 29, 30). The high levels of polysaccharides in the OL are known to be used as a bacterial decoy for antimicrobial peptides, for respiratory tract colonisation, pathogenesis, cellular invasion, antiphagocytosis (16, 30-32). The presence of high levels of polysaccharides in the TOL of rifampicin persister *M. tuberculosis* cells brings up another role for the OL polysaccharides in mycobacterial physiology. Although differences in terms of relative abundance of polysaccharides were observed, the molecular composition of OL between the MLP and the persister cells was comparable.

**Table 1.** Relative abundance of TOL^*a*^ compounds in rifampicin persister cells with respect to MLP^*b*^ cells as detected by GC-MS

| Compound | Fold increase in Persister cells<br>(w.r.t. MLP cells) |
| --- | --- |
| α-D-Glucopyranoside | 6.67 |
| 1, 2, 5-linked-Mannitol | 4.97 |
| 3, 4-linked Mannose | 2.22 |
| Hexa-acetyl-Mannitol | 1.84 |
| 1, 2, 3-Propanetriol | 1.83 |
| Methyl 1, 2, 3, 4-Tetrahydronaphthalene-2-Carboxylate | 1.76 |
| 1, 3-Di-Iso-Propylnaphthalene | 1.73 |
| 1, 2, 4-linked arabinitol | 1.68 |
| D- (1, 2-linked mannitol) | 1.58 |
| 1, 3-Di-Iso-Propylnaphthalene | 1.41 |
| 5-linked Galactonitrile | 1.30 |
| Galactose pyranoside | 1.11 |
| 1, 7-Di-Iso-Propylnaphthalene | 1.10 |
| α- D-Mannopyranoside | 1.08 |

<sup>a</sup>Thick Outer Layer; <sup>b</sup>Mid-log phase

### Persister cells restrict rifampicin permeability

Reduced cell wall permeability is known to be a factor contributing to antibiotic resistance in *Neisseria meningitidis* and *Staphylococcus aureus* (33-35). In mycobacteria also, the role of membrane permeability in rifampicin resistance in the actively growing populations of *Mycobacterium intracellulare* has been reported (36). Therefore, presuming that the TOL might restrict the entry of rifampicin, experiments were carried out to understand the permeability characteristics of rifampicin persister cells. For this purpose, 5-carboxyfluorescin (5-FAM) conjugated rifampicin (5-FAM-rifampicin) was used to monitor entry of the same into persister cells, in comparison with its entry into MLP cells.

Conjugation of the small hydrophobic fluorophore, 5-FAM, to rifampicin generated two possible isomers of 5-FAM-rifampicin depending on the ester bond formed on the two aliphatic hydroxyl group on rifampicin molecule (**Fig. 3A**). 5-FAM group was selected due to the nonpolar nature and smaller size of the fluorophore. This was to avoid any kind of alteration in the polarity of the molecule and to minimise the molecular size to maintain its entry into rifampicin-unexposed cells. Thus, it is like rifampicin, which is nonpolar in nature and believed to passively diffuse through the mycobacterial cell wall and accumulate inside the mycobacterial cell within 20 min of exposure (37). Further, the conjugation of 5-FAM to rifampicin reduced the bactericidal activity of the antibiotic to 2.5% of its original bioactivity, as calculated from the bio-assay (**Fig. 3B; Fig. S3**). An average zone of inhibition of 1.77 ± 0.035 cm was obtained for 5-FAM-rifampicin at a concentration of 29.2 µg/ml, while 0.74 µg/ml of rifampicin gave the same zone of inhibition, showing the drop in bioactivity of 5-FAM-rifampicin to 2.5% compared to that of rifampicin. Owing to these molecular properties, use of 5-FAM-rifampicin avoided inflicting lethality on the cells which would have otherwise affected the permeability assay. Radioactively labelled native rifampicin was not used as it would have affected the viability of the cells. The concentration-dependent entry of 5-FAM-rifampicin into MLP cells confirmed its high level of permeability (**Fig. 3C, D**). The extent of increase in the permeability showed a linear correlation to the concentration of 5-FAM-rifampicin (**Fig. 3E**). This indicated that the conjugation of 5-FAM to rifampicin did not affect its uptake by the actively growing MLP cells. The permeability characteristics of 5-FAM-rifampicin into MLP cells validated that the uptake of 5-FAM-rifampicin by the persister cells could be considered as a measure of its extent of permeability into the persister cells as well, in comparison to MLP cells.

**FIG. 3.**
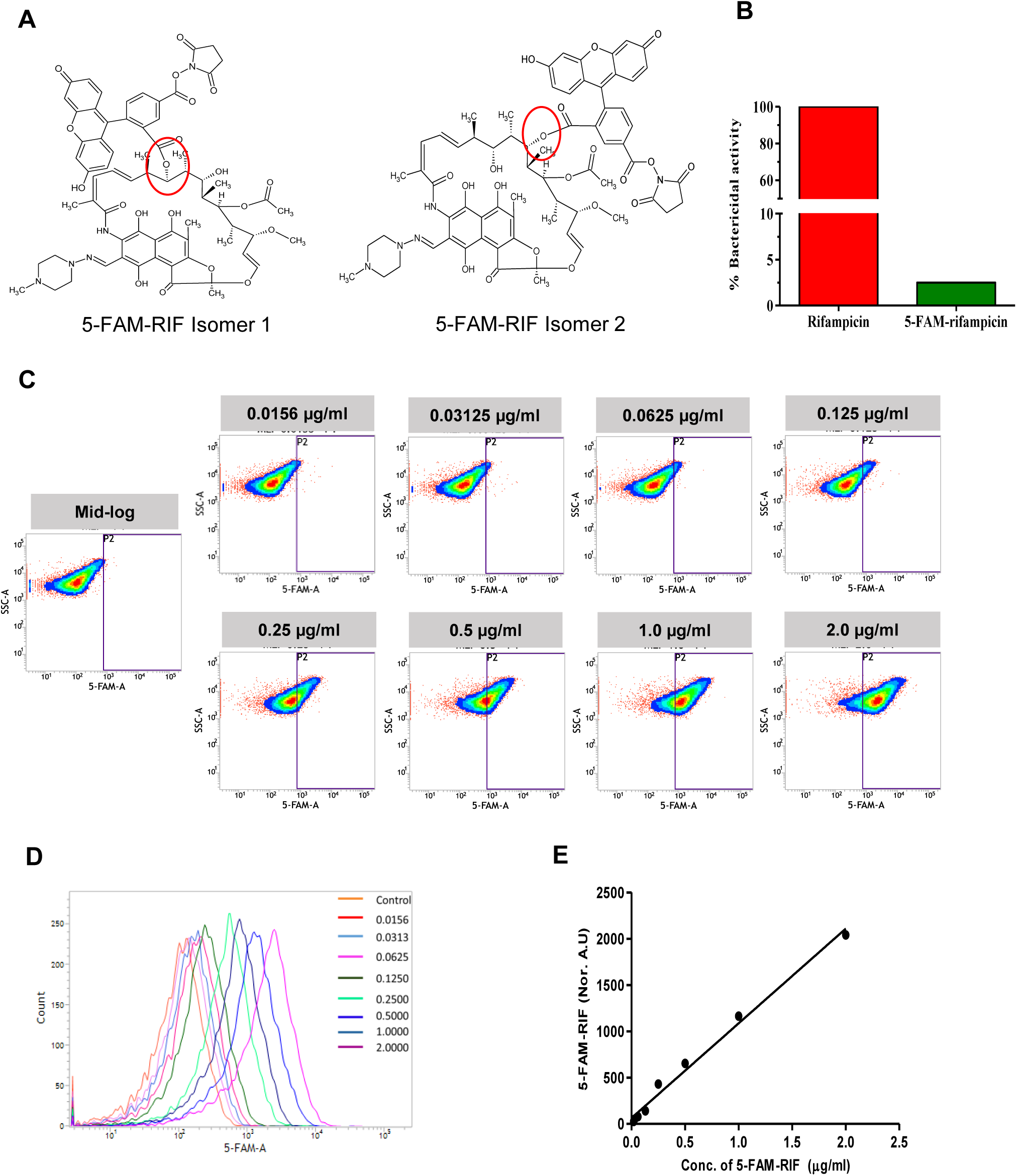
Characterisation of 5-FAM-rifampicin permeability into MLP and rifampicin persister cells. (**A**) The possible structural isomers of 5-FAM-rifampicin conjugates. The carboxyl group of the 5-FAM can form an ester bond with any one of the hydroxyl groups in the aliphatic ring of rifampicin, generating two possible isomers. Ester bond is shown in red circle. (**B**) Bar graph showing the relative bactericidal activity of rifampicin and 5-FAM-rifampicin using *Staphylococcus aureus* by agar diffusion assay. (**C**) Density plots and (**E**) histogram overlay of MLP cells incubated with increasing concentrations of 5-FAM-rifampicin by flow cytometry. (**E**) Standard graph showing the 5-FAM-rifampicin permeability into MLP cells at two-fold increasing concentrations.

MLP and persister cells were incubated with 1.5 µg/ml (concentration equivalent to 10x MBC rifampicin used in our earlier study; 12) of 5-FAM-rifampicin and analysed using fluorescence microscopy. The low levels of fluorescence in the persister cells, as compared to that in the MLP cells, indicated restricted entry of 5-FAM-rifampicin into persister cells (**Fig. 4A**). Fluorescence microscopy of persister cells, which were gently bead beaten to remove TOL and incubated with 5-FAM-rifampicin, showed enhanced levels of 5-FAM fluorescence indicating increased 5-FAM-rifampicin entry into the persister cells (**Fig. 4B**). Further, we determined the relative time-dependent entry of 5-FAM-rifampicin into MLP and persister cells, with or without bead beating. For this purpose, the cells were incubated over a period of one hour with 1.5 µg/ml of 5-FAM-rifampicin and aliquots were withdrawn every 15 min followed by flow cytometry analysis. Both the native and the bead beaten MLP cells showed a time-dependent steady increase in the 5-FAM-rifampicin fluorescence showing that the thin OL of MLP cells did not play any role in the permeability of 5-FAM-rifampicin (**Fig. 4C, E, G**). On the contrary, incubation of persister cells with 5-FAM-rifampicin did not show any noticeable fluorescence inside the cells (**Fig. 4D, F upper panels, H**). This indicated restricted permeability of the rifampicin conjugate into the cells. Whereas, the bead beaten rifampicin persisters showed a significant time-dependent increase in the fluorescence for at least upto first 30 min of incubation, followed by a level of saturation, suggesting the increased permeability of 5-FAM-rifampicin (**Fig. 4D, F lower panels, H**). Thus, the removal of the TOL by bead beating allowed permeability of 5-FAM-rifampicin into the persister cells. These experiments confirmed the substantial role of TOL to function as a ‘barrier’ to restrict permeability of rifampicin into persister cells.

**FIG 4.**
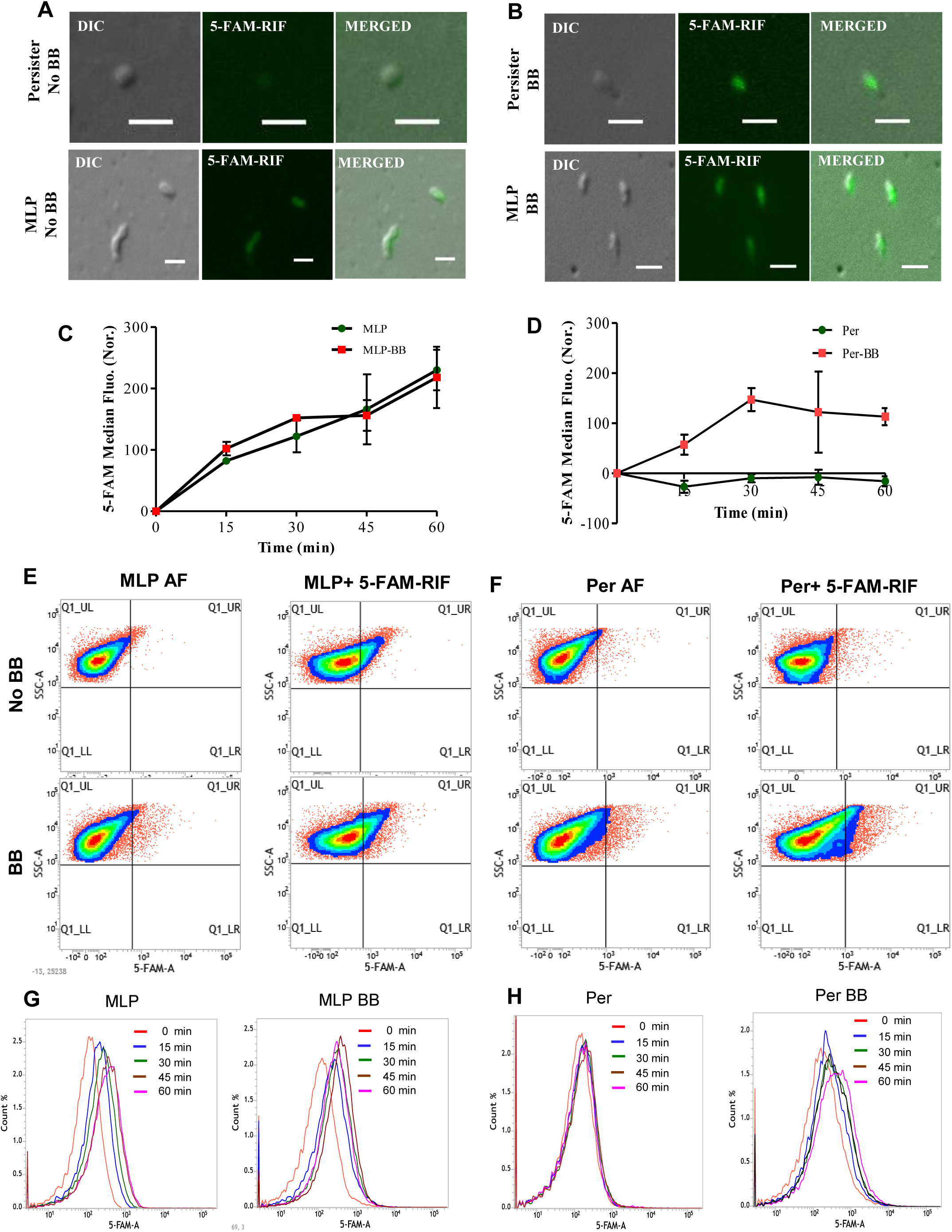
Permeability of *M. tuberculosis* to 5-FAM-rifampicin. Fluorescence microscopy images of *M. tuberculosis* rifampicin persister cells showing differential entry of 5-FAM-rifampicn in persisters (**A)** and MLP cells **(B**) before and after the removal of OL by bead beating (BB). (Scale, 2 µm). Line graph from flow cytometry analysis for the time-dependent entry of 5-FAM-rifampicin into MLP (**C**) and rifampicin persisters (**D**) with or without bead beating (BB) and its corresponding density plots (**E**) and histogram overlay (**F**). (AF, Autofluorescence).

### Verapamil did not affect the permeability of persisters to 5-FAM-rifampicin

Multidrug efflux pumps are known to contribute to antibiotic tolerance in *M. tuberculosis* (38-40). It was reported that after incubation of rifampicin-exposed *M. tuberculosis* cells in *in vitro* cultures and in infected macrophages with the efflux pump inhibitor, verapamil, rifampicin levels inside the cells increased thereby enhancing susceptibility (41, 42). This study showed that verapamil-sensitive efflux pump was involved in the removal of rifampicin from the cells. With this background information, it was of interest to find out whether efflux pumps were involved in the tolerance of *M. tuberculosis* persister cells to rifampicin *in vitro*.

For this purpose, we exposed rifampicin persister cells to 50 µg/ml of verapamil and used for 5-FAM-rifampicin permeability assay. We did not observe any difference in terms of the 5-FAM-rifampicin fluorescence intensity of verapamil-treated and untreated rifampicin persister cell samples (**Fig. 5 A, B, C**). It ruled out the possibility of rifampicin efflux as a possible mechanism for the rifampicin-tolerance in persister cells *in vitro* and further confirmed that the TOL plays a substantial role in the reduced rifampicin permeability into *M. tuberculosis* persister cells. On the contrary, verapamil was found to be an efflux inhibitor in *M. tuberculosis* infected mice, zebra fish and macrophages (40, 43). This apparent contradiction with these works is probably due to the difference in the physiological status of the cells exposed to verapamil. While the cells were in the persistence phase in our study, the cells were in the infected animal model and macrophages in the other studies (40, 43).

**FIG 5.**
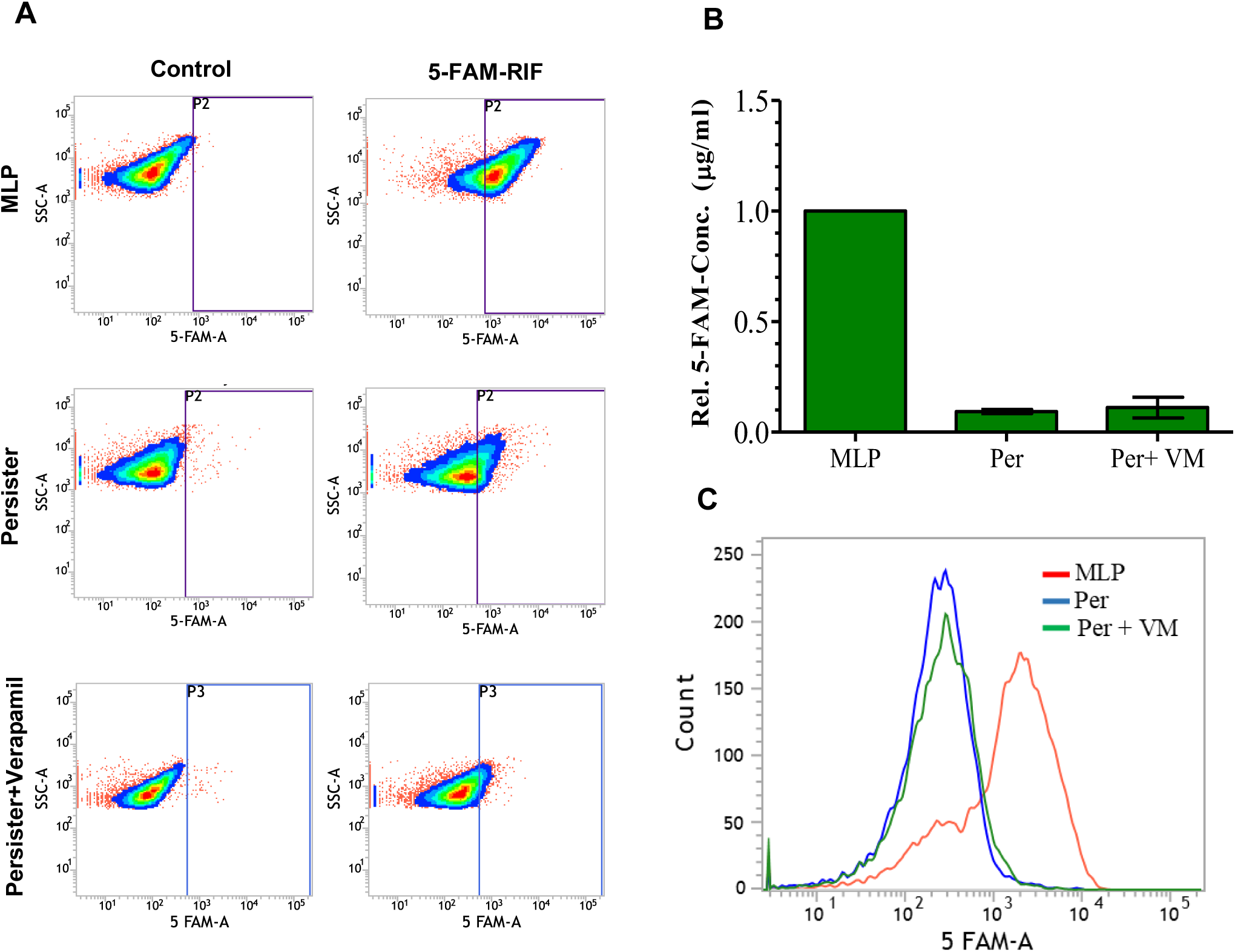
Estimation of the relative permeability of 5-FAM-rifampicin into rifampicin persisters in presence of efflux pump inhibitor verapamil. (**A**) Density plots of 5-FAM-rifampicin entry in *M. tuberculosis* cells with or without verapamil exposure. (**B**) Bar graph showing relative quantitation of the 5-FAM-rifampicin entry between MLP cells and rifampicin persisters with and without verapamil (VM) treatment. (**C**) Histogram overlay of the flow cytometry profile.

## Discussion

*M. tuberculosis* is well known for its drug tolerance and natural resistance to many antibiotics. The unique and complex cell wall features of mycobacteria imposing limited permeability have been attributed to be the reason for its natural antibiotic tolerance. Owing to the peripheral location, OL plays a significant role in macrophage interaction, cell adhesion and pathogenicity of *M. tuberculosis* (44-46). Further, metabolic and detoxifying enzymes, including penicillinase, urease, phosphatases and superoxide dismutase, are present in the OL (30). The presence of immunomodulatory factors in the OL of pathogenic *M. tuberculosis* plays a major role in the initial host immune response (47). Previous reports have shown that the OL reduces bacterial interaction with the macrophage in the absence of serum opsonins thereby possess anti-phagocytic activity in *M. tuberculosis* (16) and in Gram-negative bacteria (31, 48). Also, it has been reported that the capsular polysaccharides prevent the entry of potentially harmful host-derived macromolecules like bactericidal peptides in Gram-negative bacteria (49-51). Thus, in addition to the contribution of OL to various physiological aspects of mycobacteria (16, 29; 44-47), the present findings show yet another role for OL in the rifampicin tolerance by persisters.

The strategy seemed to be to increase the levels of polysaccharide components of OL significantly and thereby confer higher negative charge to restrict entry of rifampicin. The negative surface charge density was found varying, probably depending upon the extent of casing of the cells by the OL. The gradual increase in the surface charge density of the persister cells might be due to the accumulation of OL components over the cell surface during rifampicin exposure. The decrease in the negative surface charge density during regrowth phase indicated the specific role of OL thickening during persistence phase as a survival strategy during antibiotic exposure which is no more present on (required for) the bacteria as they have gained resistance and entered regrowth phase. Moreover, the multiple peaks of ZP values in the persister and regrowth phase cells denote the presence of heterogeneous sub-populations with different surface charge density depending on the extent of OL thickening of the cells.

It is possible that the persisters are getting benefited by the ‘barrier’ effect of TOL in limiting the entry of antibiotics. Restricted antibiotic uptake through outer membrane modification is a known mechanism of rifampicin resistance in the actively growing *M. intracellulare* and *N. meningitidis* (33, 36). Previous study has reported that the use of Tween 80 could improve the permeability of rifampicin in *M. intracellulare* in the growth medium as it is known to reduce permeability barriers in mycobacteria (36) However, in all our experiments we used 0.05% of Tween-80 containing 7H9 broth. Though it is likely that the degradation and assimilation of Tween-80 by the mycobacterial cells over the course of growth and drug exposure (52) could affect the effective concentration of the Tween-80 and thereby altered permeability, the differential OL thickness between the persisters and 12 day old rifampicin unexposed control cells exclude this possibility (see **Fig. S1**). Since the surface charge of persister cells was negative, the polar and charged nature of the OL need to be considered for its effect on the permeability of a nonpolar antibiotic such as rifampicin. The higher negative charge (polar nature) may be expected to reduce the permeability of a more non-polar type of molecule, such as 5-FAM-rifampicin. However, an integrated study using Wayne’s *in vitro* hypoxia model at pH 5.8 has shown that while many lipophilic drugs (rifampicin, rifapentine, bedaquiline, clofazimine, nitrazoxamide) could reduce cfu of hypoxic cells by ≥2-log_10_, many hydrophilic drugs (metronidazole, moxifloxacin, pyrazinamide, ethambutol, isoniazid, meropenem) could not effectively reduce the cfu of hypoxic cells (53). Therefore, the polar nature of the TOL cannot be taken as the sole reason for the restricted entry of non-polar (lipophilic) rifampicin. A combination of increased physical thickness and negative charge may be contributing to the restricted entry of rifampicin into *M. tuberculosis* persister cells, thereby helping survival and subsequent acquisition of drug resistance.

## Acknowledgements

PA dedicates this work as a tribute to Prof. T. Ramakrishnan (late), who led the pioneering and foundation-laying work on the biochemistry and molecular biology of *Mycobacterium tuberculosis* at Indian Institute of Science, Bangalore.

## Funding

The work was supported by funds from the DBT-IISc partnership programme and IISc. Authors acknowledge DBT-supported FACS facility in the Biological Sciences Division, and the infrastructure facilities supported by DST-FIST, UGC-CAS, ICMR-CAS, and IISc, in the MCB Dep’t. JS and SS received SRF from DBT and CSIR, respectively.

## Conflicts of interest statement

None declared.

## Author Contributions

PA, JS conceived/designed expts; JS, SS performed expts; PA, JS, SS analysed data; PA contributed reagents, materials, and analysis tools; PA, JS wrote the manuscript.

